# Administration of orexin A in the posterior paraventricular nucleus of the thalamus promotes cocaine-seeking behavior and is associated with hypothalamic activation

**DOI:** 10.1101/185538

**Authors:** Alessandra Matzeu, Rémi Martin-Fardon

## Abstract

Hypothalamic orexin (Orx) neurons that project to the paraventricular nucleus of the thalamus (PVT) have received growing interest because of their role in drug-seeking behavior. When injected in the posterior PVT (pPVT), OrxA reinstated extinguished cocaine-seeking behavior in rats that had long access (LgA) to cocaine for 6 h/day after an intermediate period of abstinence (I-Abst, 2-3 weeks). Considering the long-lasting nature of drug-seeking behavior and that the PVT sends projections to the hypothalamus, the present study examined whether (*i*) OrxA’s priming effect is preserved after a period of protracted abstinence (P-Abst, 4-5 weeks) in LgA rats and (*ii*) the neural activation pattern (i.e., Fos^+^ and Fos^+^/Orx^+^ cells) in the lateral hypothalamus (LH), dorsomedial hypothalamus (DMH), and perifornical area (PFA) following intra-pPVT OrxA administration that may explain OrxA-induced reinstatement in LgA animals. As reported previously, OrxA administration in the pPVT triggered cocaine-seeking behavior after I-Abst. With P-Abst, the priming effect of OrxA was absent. An intra-pPVT injection of OrxA produced a strong increase in neuronal activation (i.e., Fos expression) in the LH/DMH/PFA at I-Abst but not at P-Abst. The analysis of the activation (Fos^+^) of Orx neurons (Orx^+^) revealed an increase in Fos^+^/Orx^+^ expression in the LH/DMH/PFA at I-Abst only, thus paralleling the behavioral data. These data indicate that shortly after abstinence, PVT↔LH/DMH/PFA connections are strongly recruited in animals with a history of cocaine dependence. The lack of effect at P-Abst suggests that the function of Orx receptors and connectivity of the PVT↔LH/DMH/PFA circuit undergo significant neuroadaptations following P-Abst.

**SIGNIFICANCE STATEMENT:** A better understanding of the pathophysiological changes associated with cocaine addiction is needed to develop efficient pharmacotherapies. The paraventricular nucleus of the thalamus (PVT) and orexin (Orx) transmission within the PVT have been implicated in maladaptive (compulsive) behavior that is characteristic of drug addiction. The present study shows OrxA injections in the posterior PVT reinstates cocaine-seeking behavior in animals with a history of cocaine dependence, and this effect disappears after protracted abstinence, paralleled by the neuronal activation pattern in the hypothalamus. In subjects with a history of cocaine dependence, the function of Orx receptors and connectivity of the PVT↔ LH/DMH/PFA circuit undergo significant neuroadaptations.

## INTRODUCTION

The paraventricular nucleus of the thalamus (PVT) has attracted interest because of its connections with limbic and cortical structures that are part of the neurocircuitry that mediates drug-seeking behavior (Everitt et al., 2001; McFarland and Kalivas, 2001; Ito et al., 2002; Kalivas and Volkow, 2005; Belin and Everitt, 2008; Steketee and Kalivas, 2011). The PVT is selectively recruited during cocaine-seeking behavior that is induced by the presentation of cocaine-predictive stimuli (Matzeu et al., 2017). The integrity of the PVT is necessary for behavior that is motivated by the presentation of cocaine-predictive environmental stimuli (Matzeu et al., 2015). Some of the pivotal components of the neurocircuitry of addiction (Koob and Volkow, 2010) receive projections from the PVT, mostly from the posterior part (pPVT; Kirouac, 2015), highlighting the potential importance of this thalamic nucleus in the regulation of compulsive drug seeking that characterizes addiction. Among the wide projections that originate from the PVT, particularly interesting are the ones directed to the hypothalamus (Vertes and Hoover, 2008; Li and Kirouac, 2012; Kirouac, 2015); projections by which the PVT could influence behavior.

Orexin (Orx), also known as hypocretin, expression is restricted to a small group of neurons in the hypothalamus: lateral hypothalamus (LH), dorsomedial hypothalamus (DMH), and perifornical area (PFA; de Lecea et al., 1998; Peyron et al., 1998; Sakurai et al., 1998). Although Orx-containing neurons represent a relatively small proportion of cells, their projections are widely distributed throughout the brain (Peyron et al., 1998), suggesting that they play diverse roles in physiological functions, including energy homeostasis, arousal, sleep/wake cycles (Sutcliffe and de Lecea, 2000; Mieda and Yanagisawa, 2002; de Lecea, 2012), and the modulation of reward function (e.g., drug-seeking behavior; Harris et al., 2005; Dayas et al., 2008; Martin-Fardon et al., 2010; Jupp et al., 2011; Sakurai and Mieda, 2011; Martin-Fardon et al., 2016). Importantly, Orx neurons project to structures that control behavior that is motivated by drugs of abuse, such as septal nuclei, the central nucleus of the amygdala, the ventral tegmental area, the medial prefrontal cortex, the nucleus accumbens shell, and the PVT (Peyron et al., 1998; Baldo et al., 2003). Importantly, OrxA in the pPVT has been directly implicated in cocaine-seeking behavior (Matzeu et al., 2016), in which microinjection of OrxA directly in the pPVT reinstated (primed) extinguished cocaine-seeking behavior in animals that had a history of long access (LgA) to cocaine for 6 h/day, an established animal model of cocaine dependence (Matzeu et al., 2016).

Because of the remarkable resistance to extinction and long-term persistence of the motivation to administer cocaine in subjects with a history of cocaine self-administration (Martin-Fardon et al., 2016) and the strong recruitment of the Orx system during cocaine-seeking behavior (Martin-Fardon et al., 2016), the aim of the present study was to test the ability of microinjections of OrxA directly in the pPVT to reinstate extinguished cocaine-seeking behavior at 2-3 weeks of abstinence (i.e., intermediate abstinence [I-Abst]) or at 4-5 weeks of abstinence (protracted abstinence [P-Abst]). The pPVT sends projections back to the hypothalamus. Behavioral specialization is observed among specific hypothalamic subregions. The LH plays a role in the promotion (reinstatement) of drug seeking (e.g., Marchant et al., 2009), and the DMH/PFA plays a major role in the inhibition of this behavior (Marchant et al., 2012). A secondary aim of the present study was to compare neuronal activation in the LH/DMH/PFA (reflected by Fos expression) and Orx cell activation (reflected by dual Fos/Orx immunocytochemistry) at the two abstinence points (I-Abst and P-Abst).

## MATERIALS AND METHODS

### Rats

Forty male Wistar rats (Charles River, Wilmington, MA), weighing 200-225 g and 2 months old upon arrival in the laboratory, were housed two per cage in a temperature-and humidity-controlled vivarium on a reverse 12 h/12 h light/dark cycle with *ad libitum* access to food and water. All the procedures were conducted during the dark phase cycle and in strict adherence to the National Institutes of Health *Guide for the Care and Use of Laboratory Animals* and were approved by the Institutional Animal Care and Use Committee of The Scripps Research Institute.

### Self-administration and extinction training (Fig. 1)

Rats that were designated for cocaine self-administration were surgically prepared with jugular catheters 7-10 days before the beginning cocaine self-administration training in 6 h long-access (LgA) sessions. Each session was initiated by the extension of two retractable levers into the operant conditioning chambers (29 cm × 24 cm × 19.5 cm; Med Associates, St. Albans, VT, USA). Responses at the active lever were reinforced on a fixed-ratio 1 (FR1) schedule by intravenous (IV) cocaine (National Institute on Drug Abuse, Bethesda, MD, USA; 0.25 mg/0.1 ml/infusion), dissolved in 0.9% sodium chloride (Hospira, Lake Forest, IL, USA) and infused over 4 s. Each reinforced response was followed by a 20-s timeout (TO20) period that was signaled by illumination of a cue light above the active lever. Responses at the inactive lever were recorded but had no scheduled consequences.

**Fig. 1.**
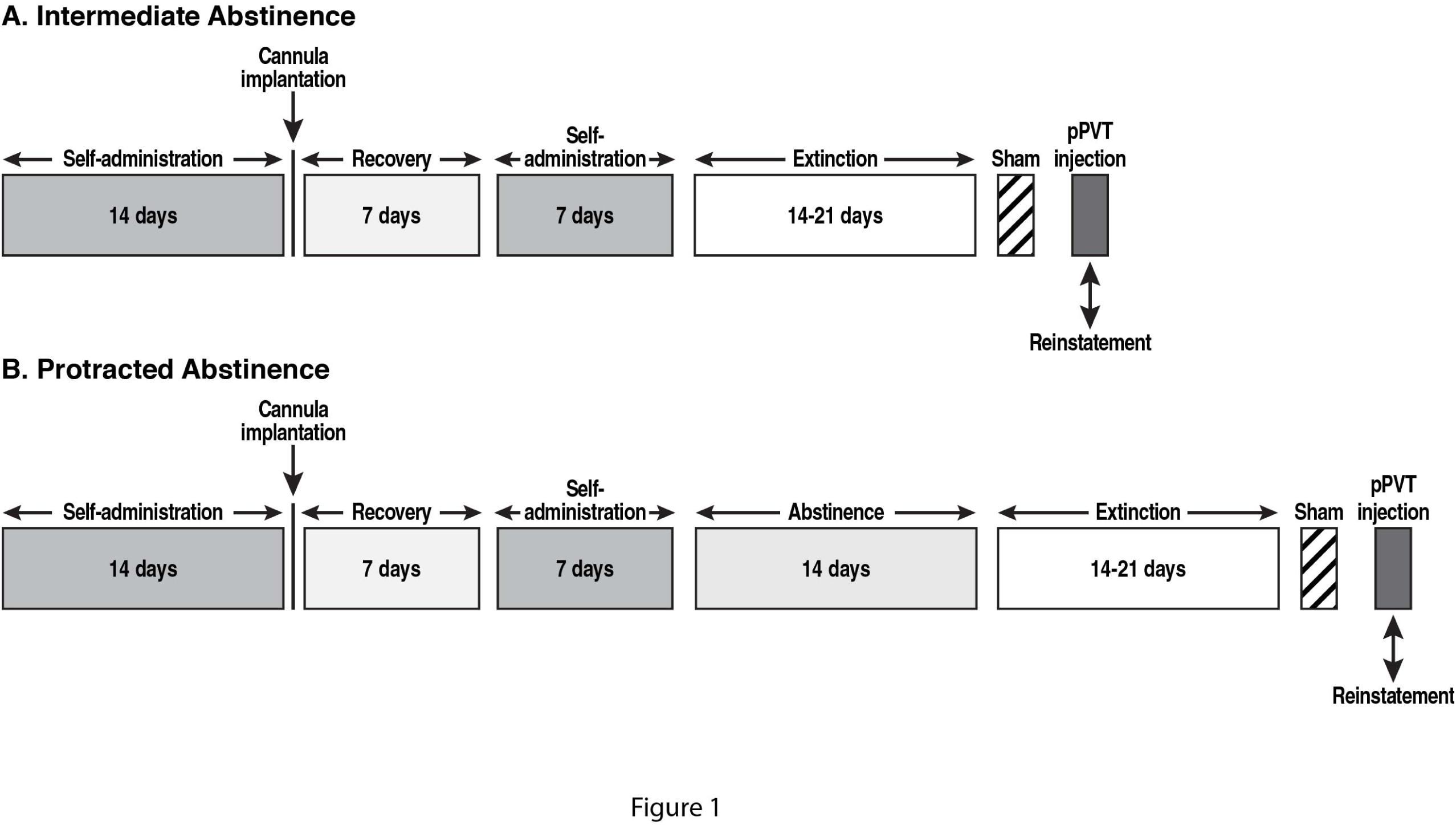
Behavioral procedure. (**A**) Intermediate abstinence. (**B**) Protracted abstinence.

#### Cannulation

Fourteen days after beginning self-administration training, the rats were implanted with a guide cannula (23-gauge, 15 mm, Plastics One, Roanoke, VA, USA) that was aimed at the pPVT (anterior/posterior, −3.3 mm; medial/lateral, ±2.72 mm from bregma; dorsal/ventral, −2.96 mm from dura, 25**°** angle; Paxinos and Watson, 1997) and positioned 3.5 mm above the target injection point (Fig. 2A, B). After 7 days of recovery, the animals resumed self-administration training for an additional 7 days.

**Fig. 2.**
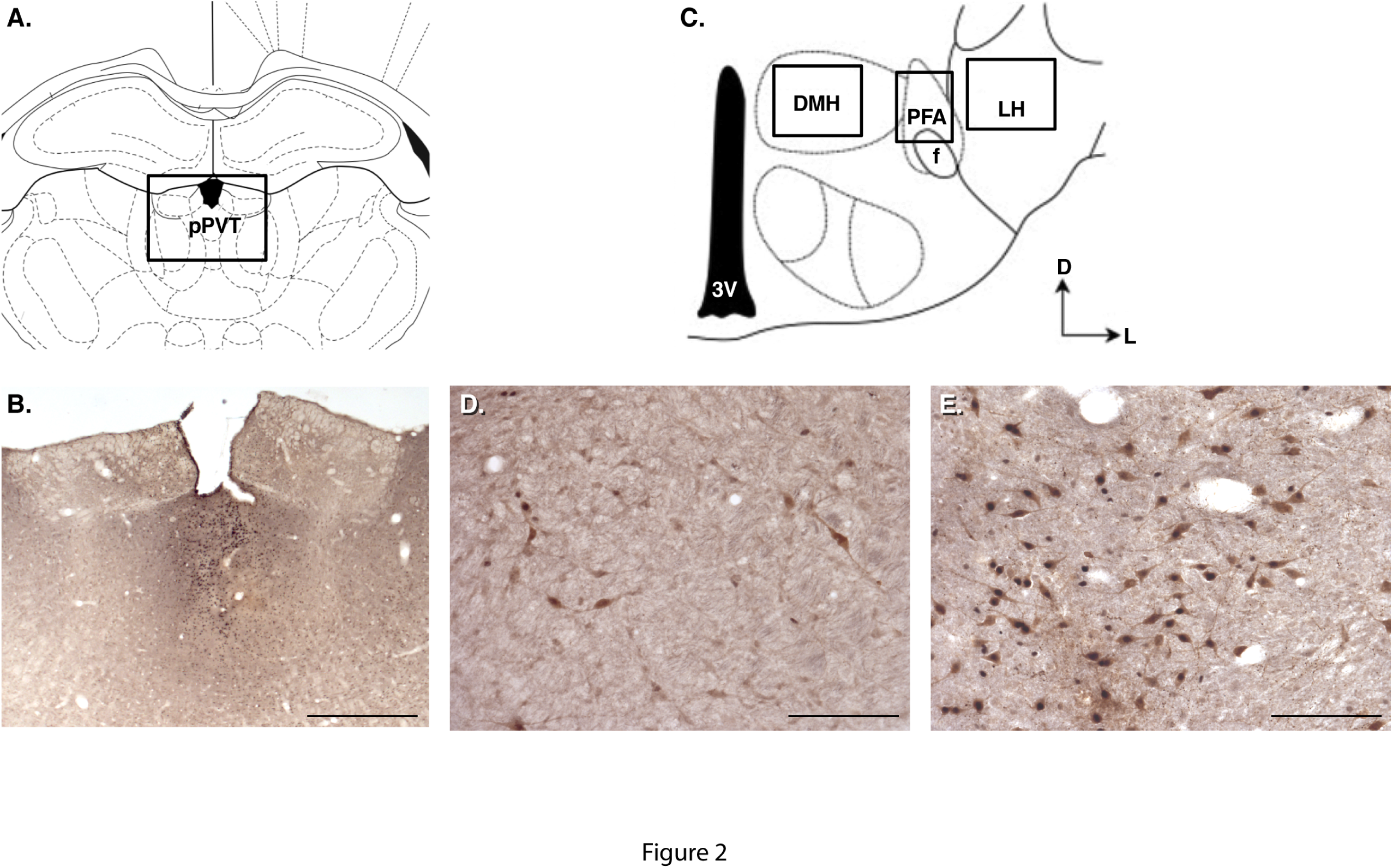
(**A**) Schematic illustration of the rostrocaudal level of the microinjection in the posterior paraventricular nucleus of the thalamus (pPVT). (**B**) Photomicrograph of injection sights (2.5× magnification) **C**. Schematic illustration of the rostrocaudal level at which Fos^+^ and Fos^+^/Orx^+^ cells were counted. LH lateral hypothalamus; DMH, dorsomedial hypothalamus; PFA, perifornical area. (**D, E**) Photomicrograph of Fos^+^ and Fos^+^/Orx^+^ LH cells in (**D**) naïve rats and (**E**) rats that self-administered cocaine and then received an injection of Orx after intermediate abstinence. The number of Fos^+^ and Fos^+^/Orx^+^ cells was significantly higher compared with naïve animals (20× magnification). Scale bars = 2000 μm in (**B**) and 250 μm in (**D**, **E**).

#### Extinction

Rats were divided in two subgroups (based on balancing for cocaine intake during self-administration training) in order to be tested under I-Abst or P-Abst. *(I-Abst)* Immediately following completion of 21 daily self-administration sessions rats to be tested following intermediate abstinence underwent extinction (EXT) training for 14 – 21 days (until the extinction criteria of ≤ 10 responses over three consecutive sessions was reached) (Matzeu et al., 2016). ***(P-Abst)*** After completion of 21 daily self-administration sessions rats to be tested following protracted abstinence were left undisturbed (just handled daily) in the vivarium for 14 days and then underwent under EXT training for 14 – 21 days (until the extinction criteria of ? 10 responses over three consecutive sessions was reached) as for I-Abst. All of the extinction sessions lasted 2 h and began with extension of the levers into the operant chambers, with the same schedule of self-administration sessions but without reward (cocaine) delivery.

### Intra-PVT microinjections (Fig. 1)

On the last day of extinction training, each rat received a sham (SHAM) injection for habituation to the microinjection. Twenty-four hours later, they received an intra-pPVT microinjection of 0.5 μg OrxA (Matzeu et al., 2016; American Peptide, Sunnyvale, CA, USA) in 0.9% sodium chloride (Hospira, Lake Forest, IL, USA) or the respective vehicle (Matzeu et al., 2016). The microinjections in the pPVT were performed using a micro-infusion pump (Harvard 22 Syringe Pump, Holliston, MA, USA) and injectors that extended 3.5 mm beyond the guide cannula. The injections were performed at a flow rate of 0.5 μl/min over 1 min. The injectors were left in place for an additional minute to allow for diffusion away from the injector tip. Following the injections, the rats were returned to their home cages for 2 min and then placed in the operant chambers under extinction conditions for 2 h.

### Immunocytochemistry

Immediately after the reinstatement session, the rats were deeply anesthetized and transcardially perfused with cold 4% paraformaldehyde in 0.1 mM sodium tetraborate, pH 9.5. Brains were removed, postfixed in 4% paraformaldehyde overnight, and stored in 30% (w/v) sucrose, 0.1% (w/v) sodium azide, and potassium phosphate buffered saline (KPBS) solution. The brains were sectioned coronally (40 μm) on a cryostat at −20°C. The sections were then processed for Fos and OrxA immunodetection. Briefly, coronal sections were blocked for 90 min using 5% normal donkey serum, 0.1% bovine serum albumin (BSA), and 0.3% Triton-X in PBS, followed by incubation for 24 h at room temperature with anti-Fos antibody (1:1000, rabbit monoclonal antibody, Cell Signaling, Danvers, MA, USA) and anti-OrxA antibody (1:15000, goat, Santa Cruz Biotechnology, Dallas, TX, USA). The tissue sections were then incubated with ImmPress reagent with secondary antibodies for 90 min (anti-rabbit IgG and anti-goat, Vector Laboratories, Burlingame, CA, USA). Fos and OrxA immunostaining was visualized using DAB-Ni (Vectors Laboratories, Burlingame, CA, USA). Fos-positive (Fos^+^) and Fos/Orx-positive (Fos^+^/Orx^+^) cells were counted within sections that incorporated the LH, DMH, and PFA (typical range: −2.40 and −3.48 from bregma; Fig. 2C-E). For immunocytochemistry, two age-matched naïve groups of rats (one for I-Abst time point and one for P-Abst time point) were prepared. The naïve rats were never exposed to the behavioral chambers. The brain tissues were then processed as above for the other animals.

### Statistical analysis

The behavioral data (cocaine intake during self-administration) were analyzed using one-way repeated-measures analysis of variance (ANOVA). The number of days to meet the extinction criterion under I-Abst and P-Abst conditions were compared using unpaired *t*-tests. Differences in behavioral responding between the final extinction sessions (EXT), the sham extinction session (SHAM) and the reinstatement extent after vehicle injection (VEH) were analyzed using two-way ANOVA between groups (I-Abst *vs*. P-Abst). The effects of OrxA on reinstatement were analyzed using two-way ANOVA, with group (I-Abst *vs*. P-Abst) and dose (vehicle *vs*. OrxA) as factors, followed by separated one-way ANOVAs for each group (I-Abst and P-Abst). The histochemical data (number of Fos^+^ and Fos^+^/Orx^+^ cells) were analyzed using two-way ANOVA, with group (I-Abst *vs*. P-Abst) as the factor, followed by separate one-way ANOVAs for each group. Significant effects in the ANOVAs were followed by the Tukey or Dunnett *post hoc* test as appropriate.

## RESULTS

Five rats were lost (two because of health complications and three because of injection misplacement), reducing the number of animals to 35 (*n* = 6 naïve and *n* = 11 for self-administration I-Abst; *n* = 6 naïve and *n* = 12 for self-administration P-Abst).

### Cocaine self-administration training and extinction

In daily 6 h cocaine self-administration sessions, the amount (mg/kg) of cocaine intake gradually increased (repeated-measures ANOVA, *F*_22,440_ = 36.57, *p* < 0.0001) from day one to day seven and then stabilized thereafter (Fig. 3). At the end of extinction training, the rats reached a comparable 3-day average (± SEM) number of responses between I-Abst (7.5 ± 1.0 responses) and P-Abst (10.4 ± 1.2; unpaired *t*-test, *t*_21_ = 1.723, *p* = 0.0996). No differences were found in the number of sessions that were required to reach the extinction criteria between groups (I-Abst: 17.3 ± 0.3 days, P-Abst: 18.0 ± 0.2 days; unpaired *t*-test, *t*_21_ = 0.6200, *p* = 0.5519).

**Fig. 3.**
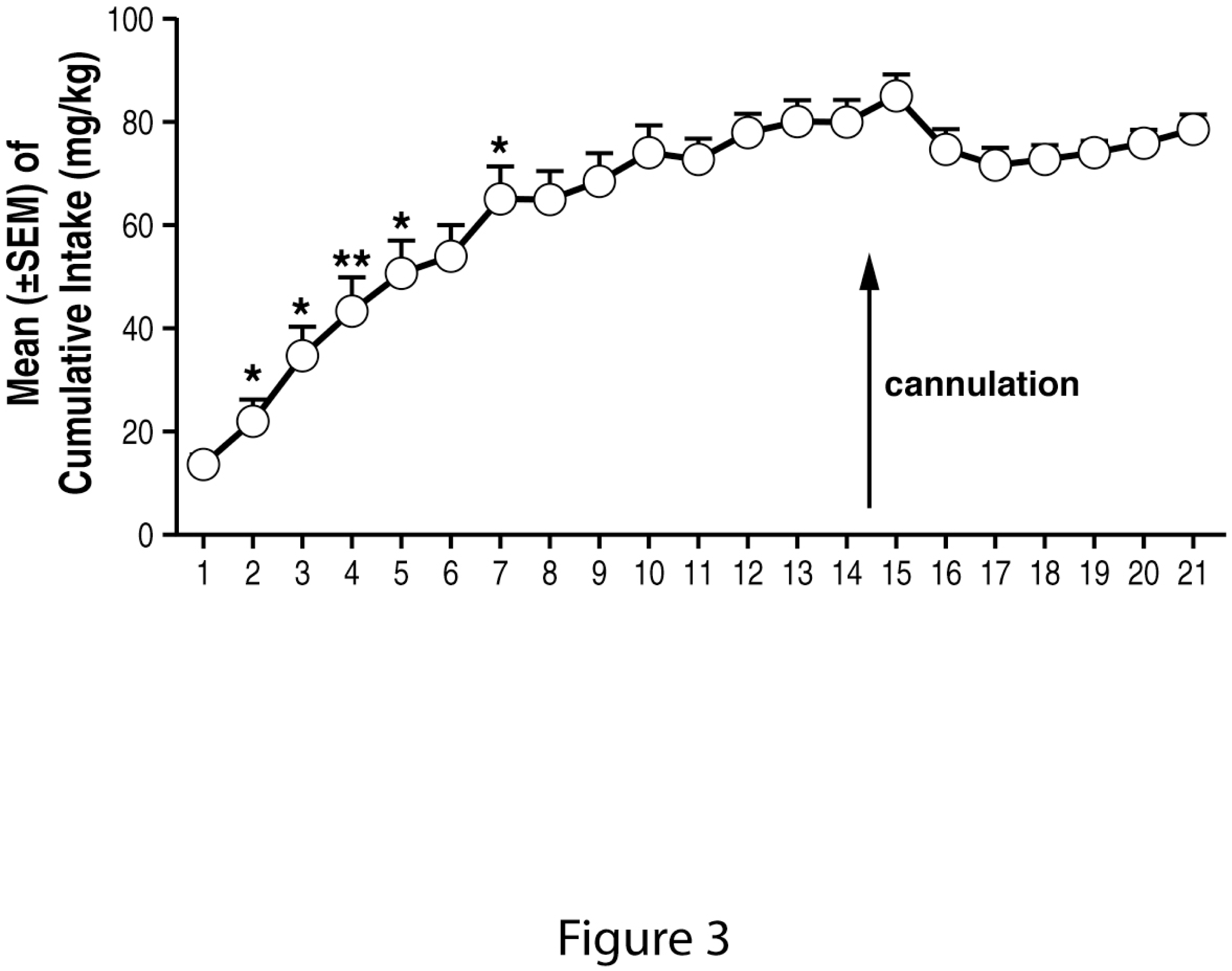
Time course of cocaine self-administration over the 21 days of training. \**p* < 0.05, \*\**p* < 0.01, *vs*. previous day (Tukey *post hoc* test). The data are expressed as mean ± SEM. *n* = 23.

### Effect of pPVT-OrxA injection

No differences were observed in the number of responses during EXT, SHAM, and VEH injections between I-Abst and P-Abst (two-way repeated-measures ANOVA; group [I-Abst, P-Abst], *F*_1,15_ = 0.6821, *p* = 0.4218; treatment [EXT, SHAM, VEH], *F*_2,15_ = 0.0597, *p* = 0.9422; group × treatment interaction, *F*_2,15_ = 0.1528, *p* = 0.8596; Fig. 4). Responses at the inactive lever did not change during these procedures (Fig. 4). Following the vehicle injection, the mean (± SEM) number of responses was negligible and not significantly different between groups (Fig. 4). The injection of OrxA in the pPVT reinstated cocaine-seeking behavior (two-way ANOVA; group [I-Abst, P-Abst], *F*_1,19_ = 4.288, *p* = 0.05; dose [VEH, ORX], *F*_1,19_ = 15.64, *p* = 0.0008; group × dose interaction, *F*_1,19_ = 5.19, *p* = 0.0344; Fig. 4). The Tukey *post hoc* test confirmed significant higher reinstatement at I-Abst *vs*. P-Abst after the OrxA injection (*p* < 0.001; Fig. 4). Dunnett’s test for each group following separate one-way ANOVAs confirmed that OrxA reinstated cocaine seeking at I-Abst (ANOVA: *F*_3,29_ = 17.26, *p* < 0.001) but not at P-Abst (*F*_3,32_ = 2.339, *p* = 0.0862; *post hoc* test, *p* < 0.001). Responses at the inactive lever were negligible and unaltered by the OrxA injection (one-way ANOVA; I-Abst, *F*_3,29_ = 0.4927, *p* = 0.6902; P-Abst, *F*_3,32_ = 2.339, *p* = 0.1029; Fig. 4).

**Fig. 4.**
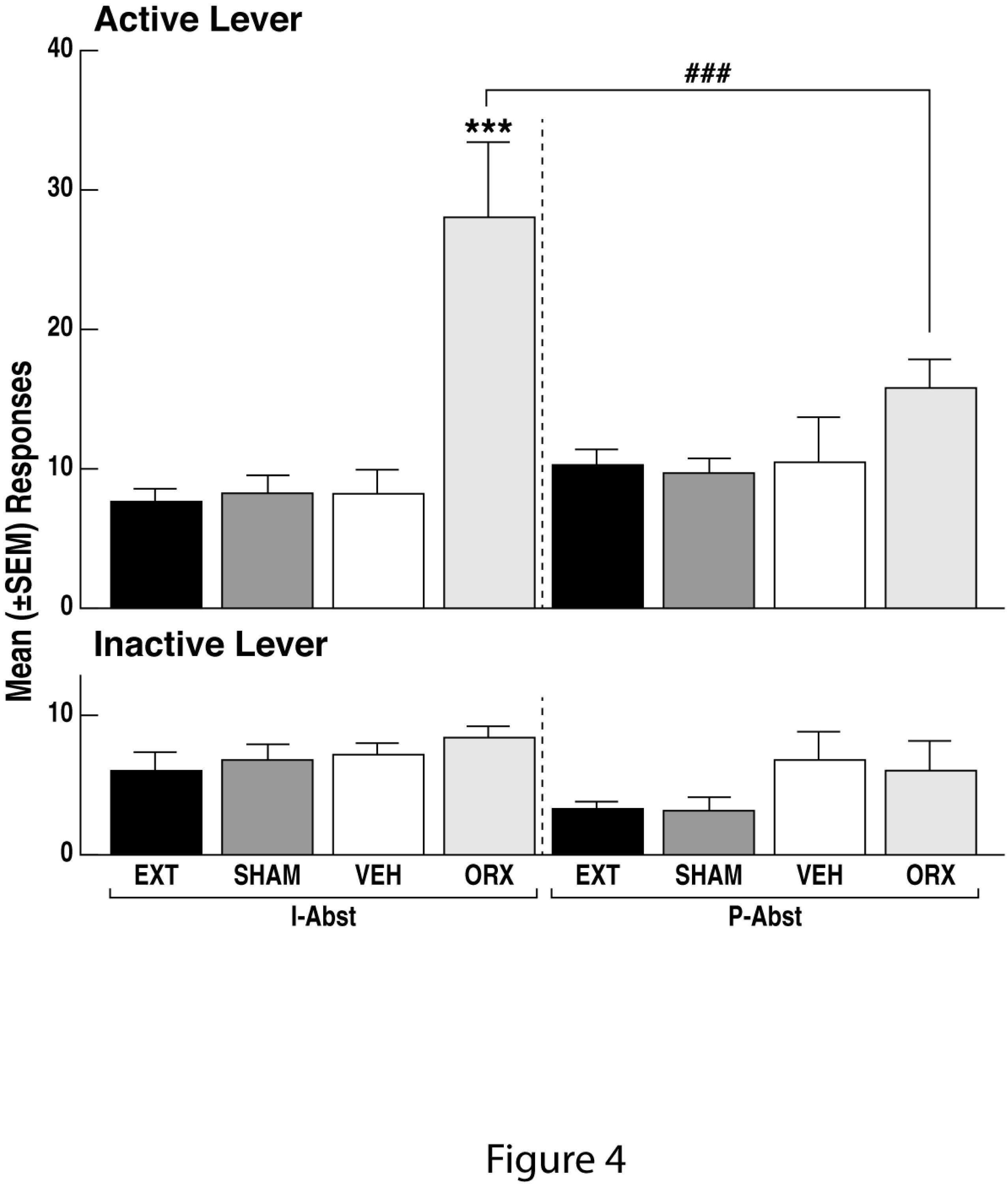
Differential reinstating effects of intra-pPVT OrxA. The animals exhibited comparable responding during extinction, sham, and vehicle injections during intermediate and protected abstinence. EXT, extinction; SHAM, sham injection; VEH, vehicle injection; ORX, orexin A. Intra-pPVT administration of OrxA induced cocaine-seeking behavior after intermediate abstinence (I-Abst) but not after protracted abstinence (P-Abst). \*\*\**p* < 0.001, *vs*. VEH; ^###^ *p* < 0.001, *vs*. corresponding protracted abstinence. The data are expressed as mean ± SEM. = 5-6 animals/group.

### Neuronal activation in the dorsal hypothalamus: LH, DMH, and PFA

Paralleling OrxA-induced reinstatement at I-Abst and the lack of reinstatement at P-Abst (Fig. 4), the intra-pPVT OrxA injection increased Fos expression at I-Abst *vs*. P-Abst in the LH (two-way ANOVA; group [I-Abst, P-Abst], *F*_1,29_ = 4.149, *p* = 0.0501; treatment [naive, VEH, ORX], *F*_2,29_ = 7.416, *p* = 0.0025; group × treatment interaction, *F*_2,29_ = 2.430, *p* = 0.1058; Fig. 5A), DMH (two-way ANOVA; group [I-Abst, P-Abst], *F*_1,29_ = 14.15, *p* = 0.0008; treatment [naive, VEH, ORX], *F*_2,29_ = 11.55, *p* = 0.0002; group × treatment interaction, *F*_2,29_ = 8.504, *p* = 0.0012; Fig. 5B), and PFA (two-way ANOVA; group [I-Abst, P-Abst], *F*_1,29_ = 1.690, *p* = 0.2038; treatment [naive, VEH, ORX], *F*_2,29_ = 13.50, *p* < 0.0001; group × treatment interaction, *F*_2,29_ = 7.771, *p* = 0.0020; Fig. 5C). Separate oneway ANOVAs confirmed that OrxA increased neuronal activation in the LH (one-way ANOVA, *F*_2,14_ = 10.70, *p* = 0.0015; *post hoc* test, *p* < 0.001), DMH (one-way ANOVA, *F*_2,14_ = 11.55, *p* = 0.0011; *post hoc* test, *p* < 0.001), and PFA (one-way ANOVA, *F*_2,14_ = 30.69, *p* < 0.001; *post hoc* test, *p* < 0.001) at I-Abst (Fig. 5) but not at P-Abst (LH: oneway ANOVA, *F*_2,15_ = 0.6112, *p* = 0.5557; DMH: one-way ANOVA, *F*_2,15_ = 0.3187, *p* = 0.7319; PFA: one-way ANOVA, *F*_2,15_ = 2.046, *p* = 0.1638; Fig. 5).

**Fig. 5.**
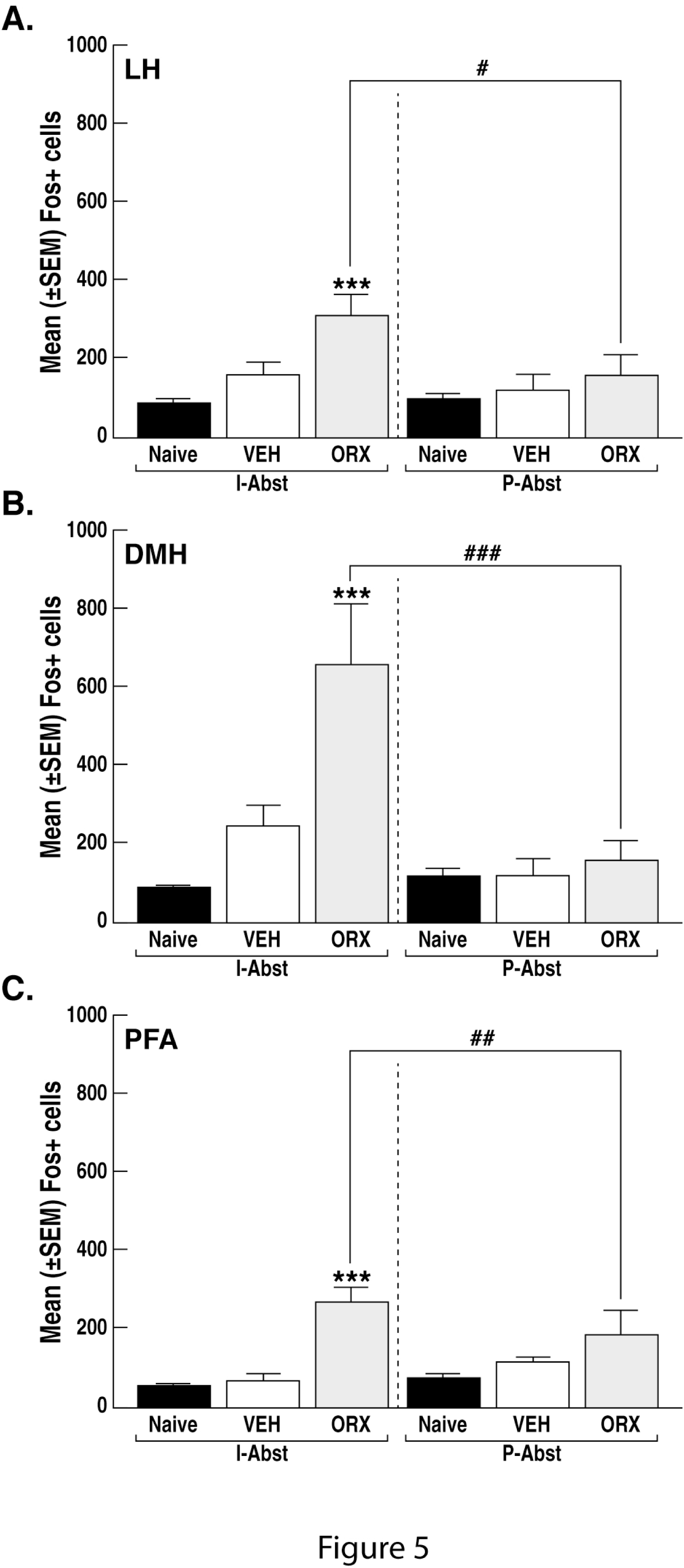
Number of Fos^+^ cells in the LH, DMH, and PFA after intermediate abstinence (I-Abst) and protracted abstinence (P-Abst). \*\*\**p* < 0.001, *vs*. naïve; ^#^*p* < 0.05, ^##^*p* < 0.01, ^###^*p* < 0.001, vs. corresponding protracted abstinence. LH, lateral hypothalamus; DMH, dorsomedial hypothalamus; PFA, perifornical area. The data are expressed as mean ± SEM. *n* = 5-6 animals/group.

### Orexin cell recruitment in the dorsal hypothalamus: LH, DMH, and PFA

An increase in Fos^+^/Orx^+^ cells was also observed in the LH (two-way ANOVA; group [I-Abst, P-Abst], *F*_1,29_ = 12.92, *p* = 0.0012; treatment [naive, VEH, ORX], *F*_2,29_ = 3.608, *p* = 0.0399; group × treatment interaction, *F*_2,29_ = 1.081, *p* = 0.3527; Fig. 6A), DMH (two-way ANOVA; group [I-Abst, P-Abst], *F*_1,29_ = 3.196, *p* = 0.0843; treatment [naive, VEH, ORX], *F*_2,29_ = 10.89, *p* = 0.0003; group × treatment interaction, *F*_2,29_ = 3.937, *p* = 0.0307; Fig. 6B), and PFA (two-way ANOVA; group [I-Abst, P-Abst], *F*_1,29_ = 3.920, *p* = 0.0473; treatment [naive, VEH, ORX], *F*_2,29_ = 4.124, *p* = 0.0265; group × treatment interaction, *F*_2,29_ = 0.9766, *p* = 0.3886; Fig. 6C) at I-Abst but not at P-Abst. Separate one-way ANOVAs confirmed that the intra-pPVT injection of OrxA induced a significant increase in Fos^+^/Orx^+^ cells in the LH (one-way ANOVA, *F*_2,14_ = 7.216, *p* = 0.0278; *post hoc* test, *p* < 0.05), DMH (one-way ANOVA, *F*_2,14_ = 5.987, *p* = 0.0499; *post hoc* test, *p* < 0.05), and PFA (one-way ANOVA, *F*_2,14_ = 13.96, *p* = 0.0005; *post hoc* test, *p* < 0.001; Fig. 6) at I-Abst but not at P-Abst (LH: one-way ANOVA, *F*_2,15_ = 0.7611, *p* = 0.4844; DMH: one-way ANOVA, *F*_2,15_ = 1.307, *p* = 0.2988; PFA: one-way ANOVA, *F*_2,15_ = 2.255, *p* = 0.1392; Fig. 6).

**Fig. 6.**
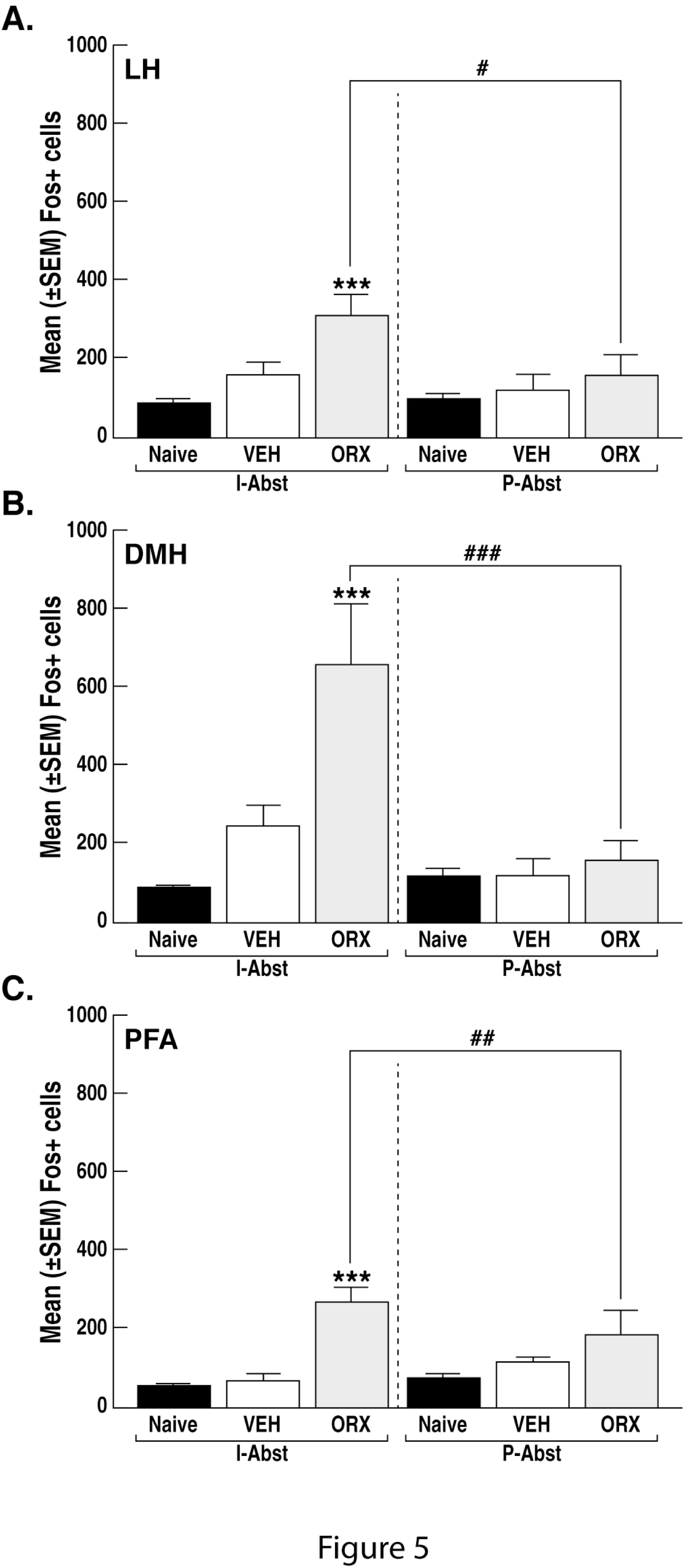
Number of Fos^+^/Orx^+^ cells in the LH, DMH and PFA after intermediate abstinence (I-Abst) and protracted abstinence (P-Abst). \*\*\**p* < 0.001, \**p* < 0.05, *vs*. naïve; ^#^*p* < 0.05, *vs*. corresponding protracted abstinence. LH, lateral hypothalamus; DMH, dorsomedial hypothalamus; PFA, perifornical area. The data are expressed as mean ± SEM. *n* = 5-6 animals/group.

## DISCUSSION

Because of the remarkable resistance to extinction and long-term persistence of cocaine-seeking behavior (Martin-Fardon et al., 2016), we tested the effects of intra-pPVT priming injections of OrxA on the reinstatement of extinguished cocaine-seeking behavior at two time points of abstinence (I-Abst and P-Abst). The pPVT sends projections to the hypothalamus, and hypothalamic subregions have different behavioral specializations. We compared Fos activation and Fos/Orx cell activation in the LH, DMH, and PFA at two abstinence time points. We found a temporal change in OrxA’s priming effects, in which strong reinstatement was observed at I-Abst, as described previously (Matzeu et al., 2016), but no priming effects were detected at P-Abst. Interestingly, the behavioral effects of OrxA were paralleled by patterns of neural activation in the LH, DMH, and PFA, reflected by strong general Fos activation in all three subregions and the recruitment of Fos^+^/Orx^+^ cells at I-Abst. These activation patterns were not observed at P-Abst. These data suggest that the functionality of Orx receptors and connectivity of the PVT↔LH/DMH/PFA circuit undergoes significant neuroadaptations following P-Abst.

Local injections of OrxA in the pPVT induced cocaine-seeking behavior at I-Abst, supporting the importance of Orx projections to the PVT in the modulation of cocaine-seeking behavior, which is consistent with a previous study (Matzeu et al., 2016), at least during the early stage of cocaine abstinence. A tentative explanation for OrxA’s priming effects that were observed at I-Abst may be the involvement of the Orx system in arousal (Sakurai et al., 2010). The expectation of food reward activates neurons that contain Orx receptors in the PVT (Choi et al., 2010). Most neurons in the PVT are sensitive to OrxA and OrxB, and the prefrontal cortex is an important target of Orx-activated PVT neurons (Ishibashi et al., 2005; Huang et al., 2006). The present results suggest that Orx inputs to the PVT facilitate cortical activation that is linked to general arousal (Sato-Suzuki et al., 2002), which could explain the reinstatement of cocaine-seeking behavior. Moreover, OrxA administration in the PVT significantly increased dopamine levels in the nucleus accumbens (Choi et al., 2012), suggesting that the PVT is a key relay for Orx’s effects on the mesolimbic dopamine system and reward-seeking behavior. Another mechanism by which Orx induces cocaine-seeking behavior could be related to the role of the PVT in mediating anxiety-and stress-like responses, which are known to precipitate drug-seeking behavior. The PVT sends projections to the dorsolateral bed nucleus of the stria terminalis and central nucleus of the amygdala. These structures contain neurons that densely express both dynorphin and corticotropin-releasing factor (CRF; Li and Kirouac, 2008). The peptides dynorphin and CRF are implicated in the expression of negative emotional states and stress responses (Davis, 1998; Heinrichs and Koob, 2004; Shirayama and Chaki, 2006; Davis et al., 2010). Both OrxA and OrxB, when injected in the PVT, produced anxiety-like behavior in rats in the open field (Li et al., 2010a) and elevated plus maze (Li et al., 2010b), suggesting that Orx may act as a stressor and thus can precipitate drug-seeking behavior.

OrxA reinstated extinguished cocaine-seeking behavior at I-Abst, but such reinstatement was absent at P-Abst. The reason for this temporal change in the behavioral effects of OrxA is unclear. One possible explanation could be that neuroplasticity that develops during cocaine self-administration may influence the Orx system (e.g., cause a fluctuation of Orx production) that projects to the pPVT and consequently induces neuroadaptive changes (e.g., changes in receptor function) in the pPVT, reflected by the change in the priming effect of OrxA at different time points of abstinence. This hypothesis is consistent with earlier findings that described the maladaptive recruitment of the Orx system by drugs of abuse, reflected by neuroadaptive changes that occurred in the ventral tegmental area (VTA; Chen et al., 2008). The self-administration of both cocaine and natural rewards induces similar transient increases in glutamatergic function in VTA dopaminergic neurons, but this enhancement of synaptic strength is persistent and resistant to extinction only in rats that self-administer cocaine (Chen et al., 2008). Interestingly, several lines of evidence suggest that the participation of the VTA in cocaine-induced neuronal and behavioral changes requires Orx inputs. For example, the activation of Orx receptor 1 in the VTA is necessary for the development of cocaine-induced locomotor sensitization (Borgland et al., 2006). OrxA-mediated *N*-methyl-D-aspartate (NMDA) receptor plasticity in the VTA increased in rats that self-administered cocaine (Borgland et al., 2009). One speculation is that a similar scenario may exist between the LH/DMH/PFA and PVT, but this possibility requires further investigation.

Interestingly and paralleling the behavioral data, significant increases in activation of the LH/DMH/PFA and the activation of Orx-producing neurons were observed at I-Abst after discrete injections of OrxA. These data suggest that recruitment of the LH, DMH, and PFA as a whole and particularly Orx neurons within these structures plays an important role in the reinstatement of extinguished cocaine-seeking behavior in rats that have a history of LgA cocaine self-administration. The strong concomitant activation of Orx neurons in the LH/DMH/PFA was unexpected when considering the reported functional dichotomy between hypothalamic subregions, in which Orx neurons in the LH participate in the regulation of reward processes, and Orx neurons in the DMH and PFA mediate responses to stressful events (Harris et al., 2005; Harris and Aston-Jones, 2006; Plaza-Zabala et al., 2010). One interpretation for these findings is that the activation of Orx neurons in the LH/DMH/PFA after the OrxA injection was attributable to a general effect of pPVT activation that increases arousal and anxiety-or stress-like behavior. The present findings may reflect the behavioral specialization among hypothalamic subregions. The LH plays a central role in promoting/reinstating drug seeking, and the DMH/PFA plays a major role in inhibiting this behavior (Marchant et al., 2009; Marchant et al., 2010; Marchant et al., 2012). Supporting this possibility, concurrent intracranial self-stimulation of the DMH and LH decreased the reinforcing actions of self-stimulation of the LH (Porrino et al., 1983). Furthermore, administration of the inhibitory peptide cocaine-and amphetamine-regulated transcript in the DMH/PFA prevented the expression of extinction in a rat model of alcoholic beer seeking (Marchant et al., 2010). Considering the evidence that the DMH/PFA regulates extinction and decreases LH activity (Porrino et al., 1983), activation of the DMH/PFA under physiological conditions may initiate the expression of extinction by inhibiting the LH (Porrino et al., 1983; Millan et al., 2011). Neuroplasticity that develops during cocaine self-administration may prevent negative feedback from DMH/PFA neurons, such that LH neurons are no longer inhibited, reflected by general activation of the LH/DMH/PFA. In the present study, the lack of a behavioral response at the P-Abst time point was paralleled by a lack of neural activation of the LH, DMH, and PFA (see Fig. 4-6), suggesting that activation of the LH/DMH/PFA following the intra-pPVT injection of OrxA was required for the reinstatement of extinguished cocaine-seeking behavior. Although the exact mechanisms and adaptations that can explain such findings are unclear, the data support the hypothesis that Orx neurons play an important role in the reinstatement of extinguished cocaine-seeking behavior that is induced by an intra-pPVT injection of OrxA. Further studies should selectively inactivate neuronal ensembles that are activated by OrxA at the I-Abst time point using the Daun02 inactivation method in c-*fos*-lacZ transgenic rats (Koya et al., 2009), followed by tests of the selective reversal of OrxA-induced cocaine-seeking behavior.

In summary, the present study found that the discrete administration of OrxA in the pPVT elicited a priming effect that reinstated cocaine-seeking behavior in dependent animals at 2-3 weeks but not 4-5 weeks of abstinence. The activation (or lack of activation) of Orx-producing neurons in the LH/DMH/PFA paralleled OrxA’s priming effect (or lack thereof), suggesting that recruitment of the LH/DMH/PFA is essential for the manifestation of cocaine-seeking behavior. The connectivity of the PVT↔LH/DMH/PFA circuit appears to undergo significant neuroadaptations during abstinence. Future studies are necessary to assess the molecular changes (e.g., Orx and Orx receptor production) that occur with cocaine dependence and prolonged periods of abstinence. Such findings may reveal valuable targets for the treatment of persistent vulnerability to relapse associated with cocaine dependence.

## Acknowledgements

This is publication number 29546 from The Scripps Research Institute. Research support: NIH/NIDA DA033344; NIH/NIAAA AA024146 (R.M.F.). We thank E.R. Zamora-Martinez for technical assistance and M. Arends for assistance with preparation of the manuscript. The authors declare no competing financial interests.

